# Phenotype-dependent habitat choice is too weak to cause assortative mating between Drosophila melanogaster strains differing in light sensitivity

**DOI:** 10.1101/2020.05.27.119271

**Authors:** Juan Ramón Peralta-Rincón, Fatima Zohra Aoulad, Antonio Prado, Pim Edelaar

**Affiliations:** Department of Molecular Biology and Biochemical Engineering, Universidad Pablo de Olavide, Seville, Spain; Department of Physiology, Anatomy and Cell Biology, Universidad Pablo de Olavide, Seville, Spain

## Abstract

Over the last few years, matching habitat choice has gained attention as a mechanism for maintaining biodiversity and driving speciation. It revolves around the idea that individuals select the habitat in which they perceive to obtain greater fitness after a prior evaluation of their local performance across heterogeneous environments. This results in individuals with similar ecological traits converging to the same patches, and hence could indirectly cause assortative mating when mating occurs in those patches.

White-eyed mutants of *Drosophila* fruit flies have a series of disadvantages compared to wild type flies, including a poorer performance under bright light. It has been previously reported that, when given a choice, wild type *Drosophila simulans* preferred a brightly lit habitat while white-eyed mutants occupied a dimly lit one. This spatial segregation allowed the eye color polymorphism to be maintained for several generations, whereas normally it is quickly replaced by the wild type.

Here we compare the habitat choice decisions of white-eyed and wild type flies in another species, *D. melanogaster*. We released groups of flies in a light gradient and recorded their departure and settlement behavior. Departure depended on sex and phenotype, but not on the light conditions of the release point. Settlement depended on sex, and on the interaction between phenotype and light conditions of the point of settlement. Nonetheless, simulations showed that this differential habitat use by the phenotypes would only cause a minimal degree of assortative mating in this species.

## Introduction

Reproductive isolation is one of the key requisites of speciation in sexual organisms: once a group of individuals within a population stops mating with the rest, they evolve independently [1]. Reproductive isolation is an extreme form of assortative mating resulting in separate gene pools, but weaker forms of assortative mating can still help to maintain genetic polymorphism [2–7]. Assortative mating and reproductive isolation might happen due to anatomical barriers [1,3,7], but it can also be caused by behavioral isolation [2–8].

Habitat choice is one such behavior which could cause assortative mating and even reproductive isolation [1–5,7,9–16]. Optimal ecological performance in heterogeneous environments is linked to the match between phenotype and environment [17]. When the environmental variability has a spatial component, habitat choice can increase this phenotype-environment match, and thereby increase fitness [9,17]. Furthermore, organisms that can evaluate an array of possible habitats and identify the environment that provides the best fit to their current phenotype, will effectively improve their ecological performance [10,18–20]. This specific, phenotype-dependent form of habitat choice is referred to as “matching habitat choice” [15,19,21]. To the extent that mating subsequently occurs in this preferred habitat, a degree of assortative mating between individuals with similar ecological traits will occur, even in the absence of sexual selection [10,12,22]. Unfortunately, few experimental tests of this possibility have been undertaken [6,14,15,21].

Taxis, the movement towards or away from a stimulus, is perhaps one the most illustrative examples of a behavior that can cause habitat choice. Subpopulations with different taxic behavior in response to the same stimulus are expected to reduce contact and acquire a certain degree of reproductive isolation in a heterogeneous habitat [11,23]. The behavior of fruit flies of the genus *Drosophila*, for instance, is heavily influenced by light [24]. There exist at least two different known phototactic responses in adults, namely “fast” and “slow” phototaxis [25–27]. Fast phototaxis occurs immediately after a fly is disturbed – by a sudden mechanical stimulus for example - and typically involves quick escape towards the light source, presumably displaying a “run for open space” response, observed in many flying insects. This response is conserved throughout *Drosophila* and is probably the most studied kind of phototaxis [28–32]. Slow phototaxis happens in the absence of such a disturbance and can be thought of as light-dependent habitat choice. Unlike fast phototaxis, the valence and intensity of slow phototaxis is quite variable among species [24,33], phenotypes [25,34] and even (in some cases genetically identical) individuals [27,35,36].

Like any habitat preference [21], taxis can be due to genetic preferences (e.g. [11,23]), a preference for a familiar environment, or because of a local assessment of a better phenotype-environment match. Eye pigmentation affects how fruit flies perceive light and thus how they might react to it [34]. White-eyed mutants, caused by a recessive loss-of-function mutation in the gene *white*, are known to have reduced visual acuity and increased sensibility to light, receiving almost 20 times more light on their photoreceptors [37] compared to wild red-eyed flies [38,39]. It is therefore reasonable to expect that preferences for a darker or brighter habitat could depend on eye pigmentation.

Jones and Probert [2] reported that, in a mixed population of *Drosophila simulans*, white-eyed mutants were outcompeted in few generations by wild type flies in both strongly and dimly lit environments. This is because white-eyed mutants have a generalized poorer biological performance [40–42]. However, when given a mixture of strong and dimly lit environments, the allele causing the white phenotype was maintained. They further showed that normal wild type flies preferred environments with much light, whereas white-eyed flies preferred dimly lit environments, and this habitat preference reduced competitive and sexual interactions. This result suggests that assortative mating and a degree of reproductive isolation occurred, consistent with the possibility that speciation could be driven by matching habitat choice [6,10,12,15,21].

In this study we test if eye color-related slow phototaxis could also result in phenotype-based spatial segregation in another species of fruit fly, *Drosophila melanogaster*, by comparing the distributions of white-eyed and wild type flies along a light gradient.

## Methods

### The testing arena

We established a light gradient by linearly arranging 5 cages with decreasing values of illuminance. Neighboring cages were connected by a 40mm diameter opening cut in each one of the wide lateral faces. The openings of the first and final cages that did not connect with any other cage, were plugged with foam stoppers that provided an oxygen inlet.

As cages we used 165 × 105 × 105 mm (∼1.5l) transparent plastic fauna cages (Savic company) with the bottom and side faces covered on the outside by a layer of black opaque adhesive vinyl. The top was covered by a rectangular piece of transparent acrylic (2 mm thick) and the openings between cages were lined with EVA foam to prevent the flies from escaping the arena (Fig 1a). A cardboard separator could be placed between two adjacent cages to block fly movement when needed. To prevent starvation and dehydration during the experiments, a petri dish with sugary water (10gl^-1^ white sugar) was placed inside each cage.

**Fig 1.**
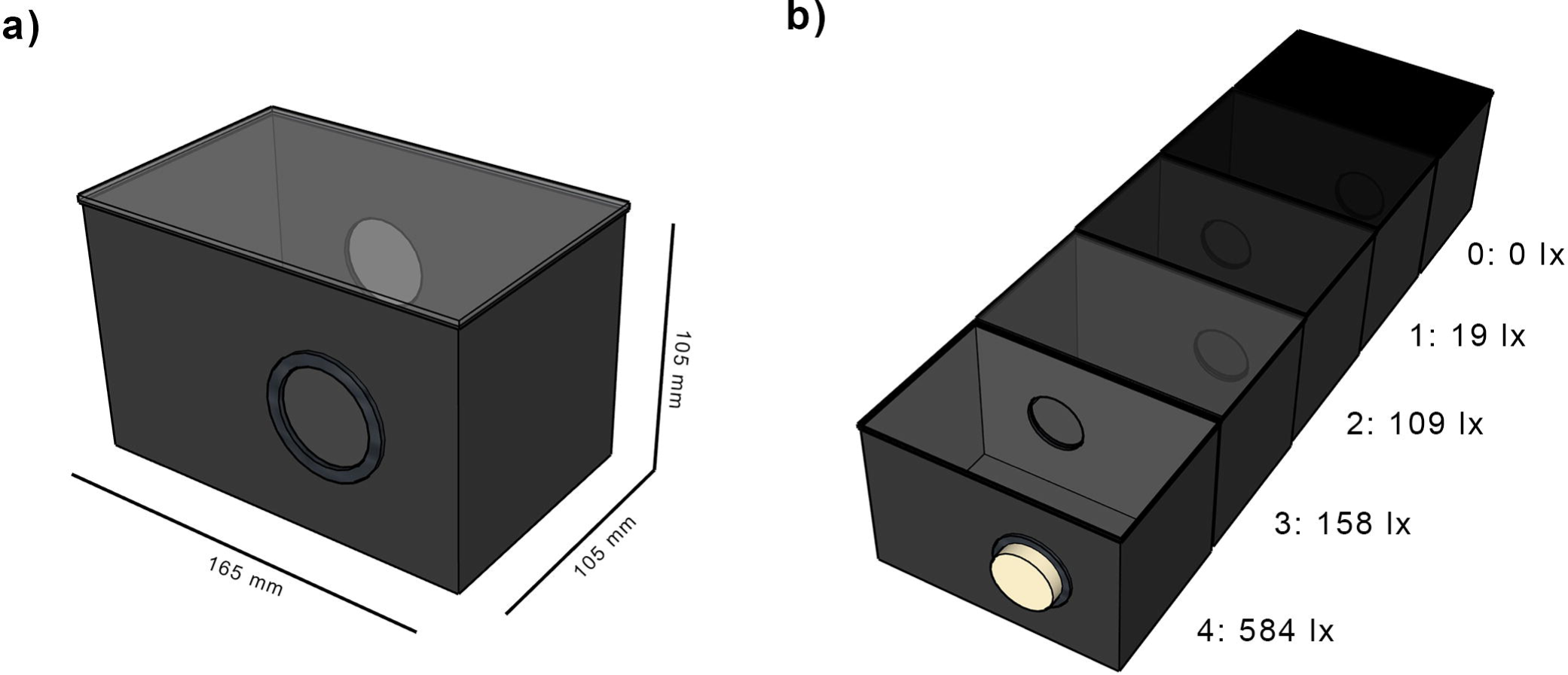
Experimental setup. Measurements for each cage module (a) and layout of all 5 cages (b). Cage 0 was covered by a layer of opaque vinyl; cages 1 to 3 were covered by 4, 2 and 1 layers of ND filter respectively. The openings on both ends of the arena were stopped by foam plugs.

The cages where lit from above by regular linear fluorescent ceiling lamps (50 Hz AC) and the light intensity gradient was achieved by covering the transparent top face of each cage with a different number of layers of Neutral Density filter paper (Lee 0.3 ND filter). For the darkest cage, we covered the top with a layer of black adhesive vinyl which prevented any light from entering. This setup created a succession of spaces with decreasing illuminance values as seen in Fig 1b.

Covering of each cage and resulting illuminance measured at the bottom of the cage with a TeckoPlus digital lux meter.

### Fly stocks

We used outbred (genetically variable) populations of wild type and *white* (w^1118^) *D. melanogaster* [43]. All the stocks were reared on conventional wheat flour and agar medium under a 12h/12h light-dark cycle at 25 °C. The experiments were performed on 2-3 days old adult flies collected from population cages, using both males and females (sex ratio close to 1). Tested flies were not reused.

### Data collection

Around 150±75 flies briefly anesthetized with CO_2_ were released in one of the cages in the morning and allowed to roam all five cages of the arena undisturbed for 20 hours (following [25]). Because dispersal decisions may depend on the initial point of release, release events were performed in each of the five cages, with two replicates per cage. In order to compare habitat choice between phenotypes, separate experiments under the same conditions were carried out with wild type *D. melanogaster* flies and white-eyed mutants, for a total of 20 release events.

After 20 hours, the openings connecting neighboring cages were blocked and the flies anesthetized again by means of CO_2_. We then counted and sexed the flies found inside each capture cage. When the cage of capture was the same as that of release, the flies were labeled as “resident”, otherwise they were labeled as “disperser”. To this data we added the information on phenotype and release cage, and an ID number for each release event.

### Data analysis

We fitted binomial Generalized Linear Mixed Models (GLMMs) [44,45] using the disperser status as the binary response variable. Sex, phenotype and either release or capture cage were used as possible predictors. For all models we added the release event ID number as a random effect in order to control for the possibility that flies were affected by unknown replicate-specific conditions. Dispersal is often divided into three separate stages: departure, transience and settlement [46], each of which might be affected by different variables. We studied the effect phenotype has on habitat choice from two different perspectives. First, using release cage as a predictor, we test whether or not the departure decision is affected by the light intensity inside the release cage. Second, using capture cage as a predictor, we test whether or not the settlement decision is determined by the intensity of the light inside the capture cage. Care must be taken when interpreting these latter results, as a high disperser/resident ratio for a specific cage can be due to increased immigration or to increased emigration. However, the combination of the two perspectives allows us to differentiate between these alternatives (see discussion).

For both of these perspectives, we fitted models with the following combinations of predictors: cage, phenotype + cage, sex + cage, sex + phenotype + cage, sex + phenotype * cage. The phenotype * cage interaction fits our main hypothesis of phenotype-dependent habitat choice, and sex is included to allow for sexual dimorphism in dispersal. We selected the model with the best fit using the Akaike Information Criterion (AIC): a lower AIC indicates a stronger support for a given model.

### Simulation of expected degree of assortative mating

Based on the obtained results on dispersal decisions (see Results), we simulated a scenario where wild and white-eyed *D. melanogaster* of both sexes are released together in an arena similar to the one described before. Our model (S1 Python Script) [47] was an individual-based simulation where each fly went through the same four steps leading to mating: release, emigration, immigration and pairing. After the final step, we determined the degree of assortative mating of the population by calculating the proportion of same-phenotype pairs out of the total number of pairs (0.5 in the absence of assortative mating, 1.0 for fully assortative mating).

In the release step, 10 flies of every possible phenotype-sex combination were assigned to each one of the 5 cages for a total of 200 flies per replicate. After that, in the emigration step, flies decided whether or not to disperse based on their sex and phenotype. During the immigration step, disperser flies were assigned to new cages based on their phenotype. The final step, pairing, involved each fly to be paired with a random individual of the opposite sex from the same cage. Each individual’s choice was independent from the rest so one mating event was recorded for each fly (except when only one sex was present inside a given cage, which was seldom the case). The probability to mate and with whom was assumed to be independent of phenotype, so we assumed no sexual selection (this might not be true in reality [40,48,49]) in order to only assess the effect of spatial distribution.

We ran a control simulation with similar parameters but in step 3 (immigration) flies were randomly assigned to their definitive cages, precluding assortative mating. We performed 1000 iterations of each simulation and compared their assortative mating scores.

## Results

The raw data (Fig 2) shows that flies tend to disperse to cage 4 (the one with most light), independent of the release cage. Nonetheless, the cage of release is always a local maximum in the distribution (Fig 2, open dots). Cage 3 seems to be avoided, except when it is the release cage.

**Fig 2.**
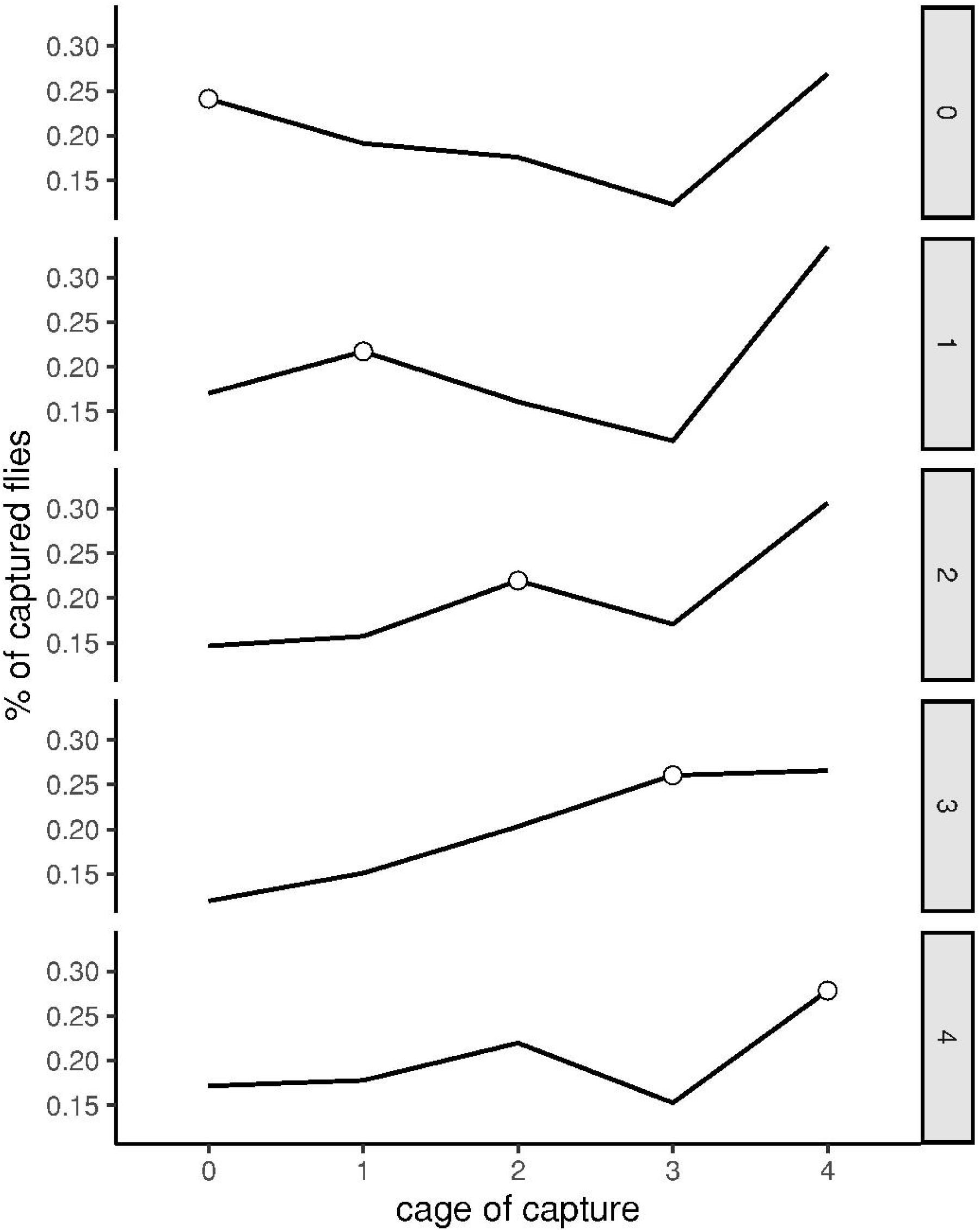
Distribution of the captured flies depending on release cage. Highlighted with an open dot is the percentage of flies that did not appear to disperse out of their release cage. Notice that the percentages are consistently high in cage 4 and low in cage 3, except when flies were released in cage 3.

### Release cage perspective

The best fitting model for the release cage approach (Table 1) suggests that the probability to disperse from the release cage depends on sex and phenotype (Fig 3), but not on the release cage itself. Starting in any given cage seems to have no effect on whether the fly departs or stays. In fact, among the fitted models, the one using release cage as the only explanatory variable fits the worst.

**Table 1.** Performance of the release cage perspective models.

| Fitted model | $\Delta AIC$ |
| --- | --- |
| **sex + phenotype** | **0.0** |
| sex + phenotype + release cage | 4.1 |
| sex | 5.1 |
| sex + release cage | 9.4 |
| sex + phenotype * release cage | 9.7 |
| phenotype | 26.0 |
| phenotype + release cage | 28.8 |
| release cage | 33.5 |

Relative performance of the fitted models compared to the best fit (in bold).

**Fig 3.**
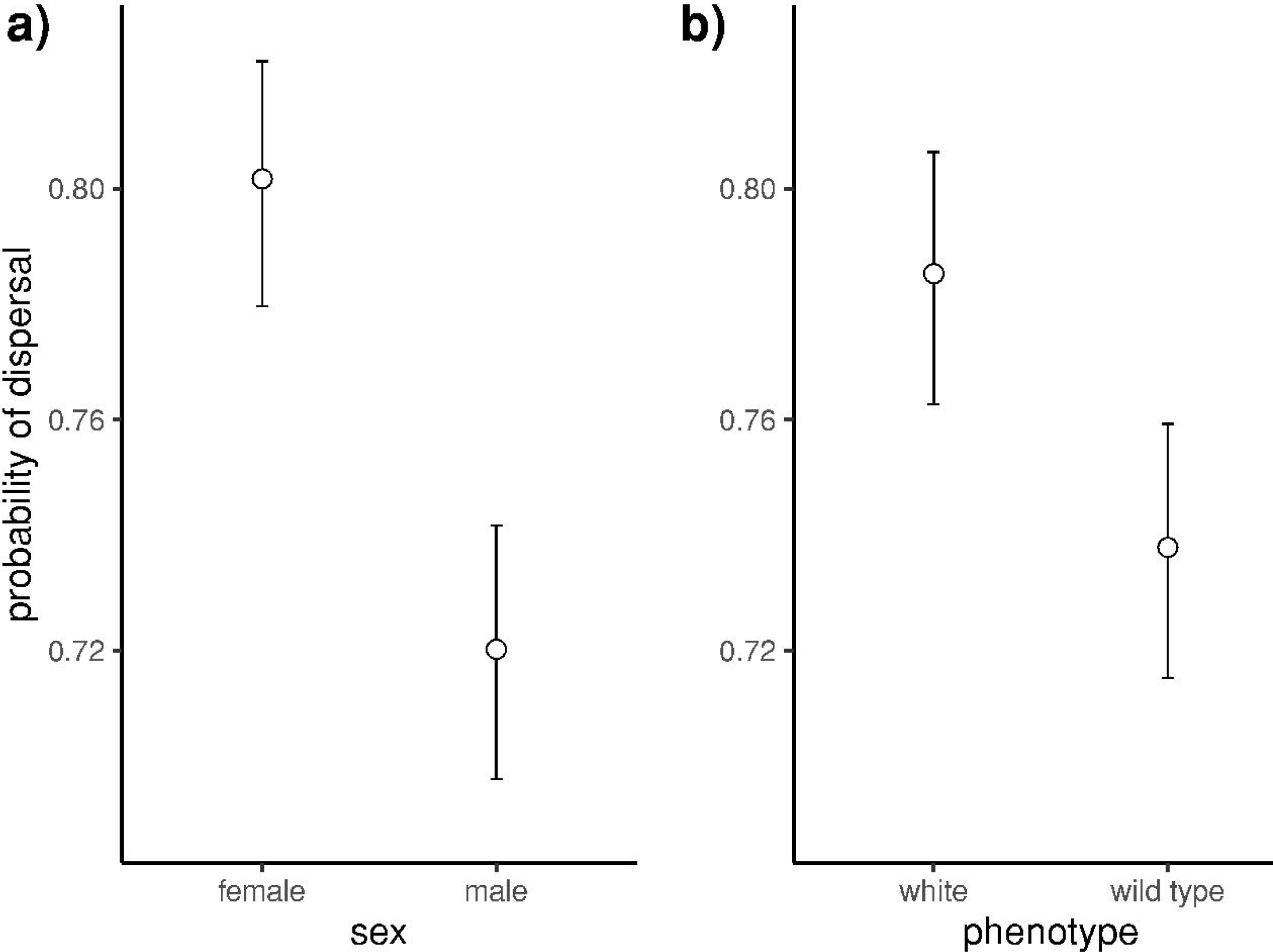
Effects of the release cage perspective model. Males are 8.1% less likely to disperse from their release cage than females (a). Wild type flies are 4.7% less likely to disperse than white-eyed mutants (b).

### Capture cage perspective

When using capture cage as a predictor, the best fitting model contains sex and the phenotype * capture cage interaction (Table 2, Fig 4). This indicates that the proportion of “resident” vs “disperser” flies captured in each cage depends on the phenotype of the flies.

**Table 2.**
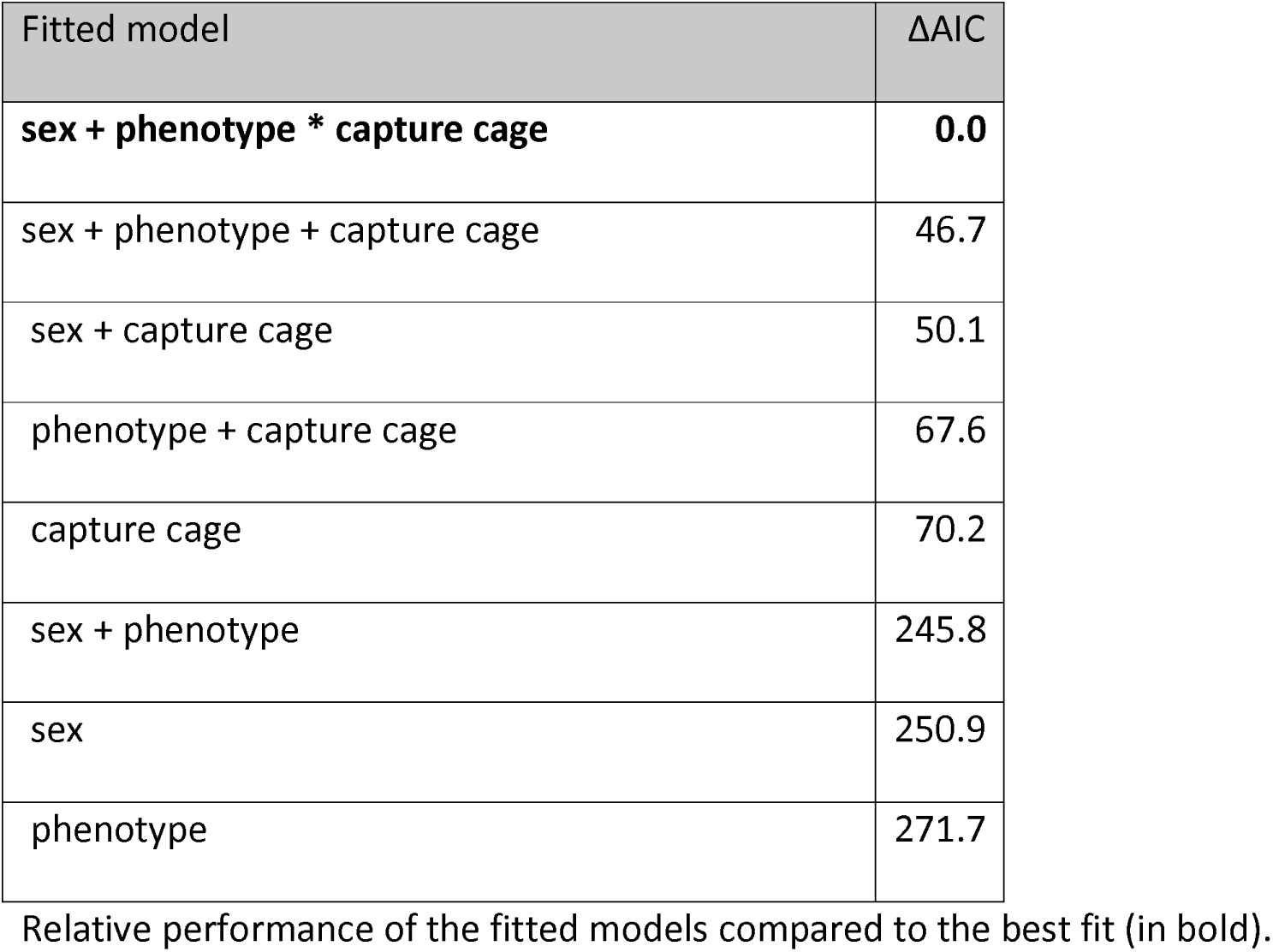
Performance of the capture cage perspective models.

| Fitted model | $\Delta AIC$ |
| --- | --- |
| **sex + phenotype \* capture cage** | **0.0** |
| sex + phenotype + capture cage | 46.7 |
| sex + capture cage | 50.1 |
| phenotype + capture cage | 67.6 |
| capture cage | 70.2 |
| sex + phenotype | 245.8 |
| sex | 250.9 |
| phenotype | 271.7 |

Relative performance of the fitted models compared to the best fit (in bold).

**Fig 4.**
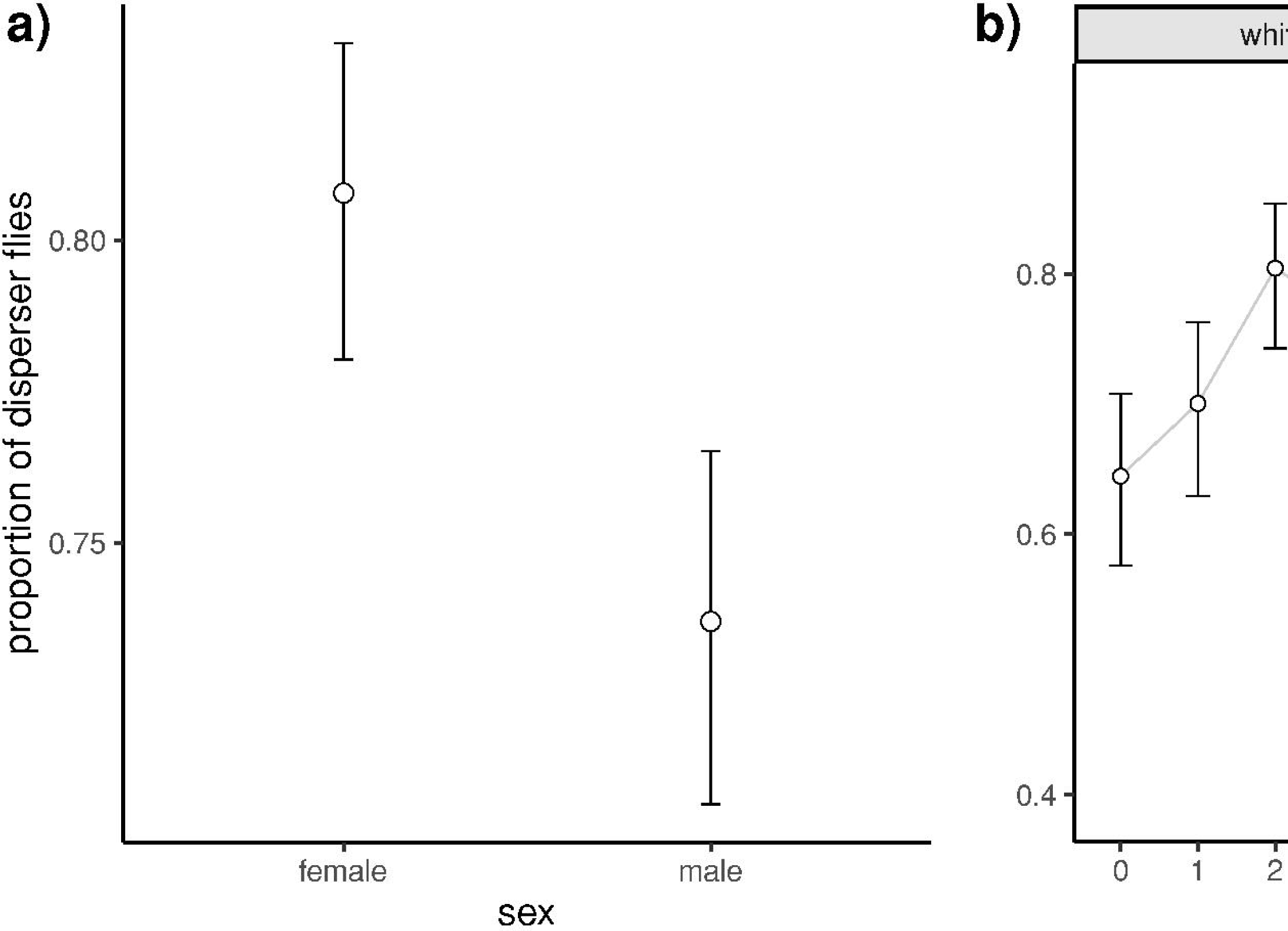
Effects of the capture cage perspective model. Effects of sex (a) and phenotype by cage of capture (b) on the proportion of disperser flies. Note for panel 4b that wild type and white-eyed disperser flies differ in their probability of choosing a particular cage. The effect is more pronounced for the wild type, but the overall patterns are rather similar (increase from cage 0 to 2, decrease for cage 3 and increase again for cage 4). In (b) an average proportion of 0.77 for all cages would be expected in the absence of habitat choice.

### Simulation results

In spite of the significant phenotype*capture cage interaction for settlement (Fig 4b), the proportion of same-phenotype pairs (0.501 ± 0.035 SD, SE=0.001) is almost the same as for when settlement is random with respect to phenotype and cage (0.500 ± 0.037 SD, SE=0.001). When the two phenotypes choose completely different capture cages for settlement, the proportion rises to 0.84; when the flies also depart for each cage based on their phenotype, this rises to 1.0 as expected.

## Discussion

The environmental gradient design of our arena is derived from the binary choice between light/dark environments described in [2] for *D. simulans*, adding a number of intermediate options. We decided on this modification after failing to obtain obvious differences in spatial preference between *D. melanogaster* phenotypes in such a binary arena, as both red and white-eyed flies clearly preferred the light environment (personal observation). This environmental gradient should provide greater resolution of phenotype-dependent habitat choice and may be closer to what flies would find in a natural situation. Indeed, we managed to detect minor differences between phenotypes with respect to settlement (Fig 4b). As a downside, it also means that accessibility to any given cage is conditioned by the release cage, but we took this into account statistically.

The fact that flies from all cages tend to disperse to cage 4 (Figs 2 and 4) can be interpreted as this cage (with the greatest amount of light) being the preferred habitat. But it is also possible that cage 4 is working as a “trap” where the bright light partially blinds and disorients the flies preventing them from exploring and choosing any of the other cages. [40] reports male white-eyed *Drosophila* having a worse courtship performance in daylight than in a dimly lit environment due to the bedazzlement caused by intense light. This coupled with poor spatial memory in white-eyed flies [50] could be biasing the distribution of the mutants towards cage 4. This is, however, less likely the case for red-eyed flies as they successfully navigate in much brighter daylight. Another possibility is that the poor food source and lack of adequate egg laying substrate enticed flies to move to bright lighting expecting a way out of the set-up, similar to what happens during fast phototaxis.

We noticed a local maximum at all release cages (Fig 2, open dots), meaning that a relatively large number of flies did not explore beyond the connection with the next cage. If this was caused by impaired movement due to handling, one would expect white-eyed flies to disperse less. This is because although biologically functional, white-eyed *Drosophila* suffer from a series of neurological disadvantages other than poor vision due to their malfunctioning *white* protein. These include reduced spatial memory [50], low copulation success [41] and progressive locomotor deficiencies [42]. For this reason, care should be taken when interpreting results from behavioral tests using these (and other) mutants, and it seems reasonable to expect that handling would negatively affect white-eyed flies more. In contrast to that, white-eyed flies were more dispersive than wild type flies (Fig 3b, Table 1), for which we do not have a good alternative hypothesis. We also found that females were more dispersive than males (Fig 3a). Potentially, searching for a suitable egg-laying site might explain this [51], in line with our above explanation of why more flies were found in cage 4.

When looking at the proportions of dispersers in capture cages, we found that it differs between the cages (Fig 4). An increase in this proportion can be caused by either high emigration (reduced number of residents) or high immigration (increased number of dispersers). Though the statistical model analyzing these proportions alone gives no clue of which process might be causing the observed results, we favor the second option because our previous results indicate that all cages have a similar probability of flies dispersing (table 1; no effect of release cage). Both phenotypes showed a photopositive response, with cage 0 being the least attractive. Both also showed a noteworthy local minimum in cage 3. If we assume the hypothesis that cage 4 is acting as a trap, this valley can be explained by cage 4 “stealing” wandering flies by preventing them from returning to cage 3. Another possible explanation is that flies cannot perceive a significant difference between cages 2 and 3 (with 109 and 158 lux respectively), which could reduce transit between those cages.

The best fitting model (table 2) also suggests that the differences in proportions of dispersers between cages is phenotype dependent: wild type flies are more selective than white-eyed mutants (Fig 4b). These findings are in agreement with [36] who linked a serotonin deficiency caused by the mutated *white* gene to high behavioral variability in white-eyed *D. melanogaster*, which would add noise to any systematic habitat choice. It is also possible that the light powered by 50Hz alternating current (used to replicate[2]) was perceived as flickering and affected white-eyed and wild type flies differently. We understand that the responsiveness of both strains to pulses decreases for frequencies above 20 Hz[39], but nonetheless it might be useful to explore the effects of DC-powered lighting in further studies.

In spite of this statistically well-supported phenotype * release cage interaction (table 2), the overall shape of the distribution is still rather similar between both phenotypes (Fig 4). While it is true that the absolute maxima are different for white-eyes and wild type flies (cages 4 and 2 respectively), these cages also attract a high proportion of flies of the other phenotype. Because of that, our simulation predicts that the observed degree of phenotype-dependent habitat choice has a minimal effect on the resulting assortative mating. Hence, if both phenotypes were released together and homogeneously across cages in this experimental set-up, we would expect only a limited amount of spatial segregation between them, and extremely little assortative mating would result. Given the competitive disadvantage of the white allele, in the long run the population would lose the allele and become homozygous for the wild type, in contrast to the results of Jones & Probert [2]. Consequently, it appears that for *D. melanogaster* phenotype-dependent habitat choice does not enable the eye color polymorphism to be maintained, as has been reported for *D. simulans*. Further experiments are necessary to determine to what extent phenotype-dependent habitat choice can maintain genetic polymorphism and even promote speciation, as has been suggested by theoretical studies [3,6,7,10,12,16,19,52] and a few empirical studies [2,53–55].

## Acknowledgments

We thank Élio Sucena, Liliana Vieira and Tânia Paulo for kindly providing the outbred fly stocks used for the research as well as useful advice on their rearing. We would also like to thank Simone Santoro and Carlos Camacho for insightful discussion regarding the analysis of our data.

## Supporting information

**S1 Python Script. Simulation of the expected degree of assortative mating.**

**S2 Dataset. Data used for the statistical analyses**

