## Supplemental_files for "Phenotype-dependent habitat choice is too weak to cause assortative mating between Drosophila melanogaster strains differing in light sensitivity": S1 Python Script.py

```
import numpy as np
import pandas as pd
import itertools

""" This program calculates D. melanogaster assortative mating due to 
habitat choice in a theoretical arena.
  
Notes:
------------
  * Sex ratio and genotype ratio = 1
  * Each fly choses ony 1 mate of the opposite sex and that pair is counted
  # females are coded as 0 and males as 1
  # white is coded as 0 and wild type as 1 

  """

# Emigration probability values
# row 0 = white-eyed phenotype
# column 0 = female

observed_emigration = np.array([[0.82, 0.75], [0.78, 0.70]])  # Observed values extracted from the "release_cage" model
forced_emigration = np.array([[1.0, 1.0], [1.0, 1.0]])        # Flies deterministically abandon their release cage
random_emigration = np.array([[0.5, 0.5], [0.5, 0.5]])        # Each fly has a 50% chance of choosing to disperse 

# Immigration probability values
# row 0 = white
# capture_cage is mapped to the columns

observed_immigration = np.array([[0.17, 0.18, 0.21, 0.20, 0.24], [0.13, 0.22, 0.25, 0.18, 0.22]])  # Observed values extracted from the "capture_cage" model
random_immigration = np.array([[0.2, 0.2, 0.2, 0.2, 0.2], [0.2, 0.2, 0.2, 0.2, 0.2]])              # Flies have equal probability of choosing any given cage
deterministic_immigration = np.array([[1.0, 0, 0, 0, 0.0], [0.0, 0.0, 0.0, 0.0, 1.0]])             # All flies with the same phenotype choose the same cage 

# Simulation functions

combinations = itertools.product((0,1), (0,1), (0,1,2,3,4))
values =  np.zeros((20, 8), dtype=np.int)
for i,c in enumerate(combinations):
    values[i, 0:3] = c

def generate_flies(n=20):
    """ Create a population of n flies with all four sex and phenotype
     combinations evenly distributed across five cages.

     Arguments
     ---------
        n: Number of flies. It will be rounded up to a multiple of 20 to ensure a perfectly balanced distribution.
    
    Returns
    -------
        pandas DataFrame
    """

    n = int(n/20)
    flies = pd.DataFrame(data = np.repeat(values, n, axis=0), columns=['sex', 'stock', 'release', 'migrate', 'capture', 'mated', 'mate', 'inbred'])
    return flies
    

def emigrate_flies(flies, emigration_probabilities=observed_emigration):
    """ Decide which of the flies disperse from the cage of release based on emigration probabilites.

     Arguments
     ---------
        flies: a DataFrame containing the output from generate_flies()

        emigration_probabilities: a 2x2 numpy array containing emigration probabilities by sex and stock
    
    Returns
    -------
        modifies the first argument in-place
    """
    
    p = [emigration_probabilities[fly[0], fly[1]] for fly in zip(flies.stock, flies.sex )] 
    flies.migrate = np.random.binomial(1, p, size=len(p))


def settle_flies(flies, immigration_probabilities=observed_immigration):
    """ Look whether or not each fly is a disperser. If yes, assign them a new cage based on 
    immigration probabilities.

     Arguments
     ---------
        flies: a DataFrame containing the output from emigrate_flies()

        immigration_probabilities: a 2x5 numpy array containing immigration probabilities by stock and cage
    
    Returns
    -------
        modifies the first argument in-place
    """

    flies.capture = [np.random.choice([0, 1, 2, 3, 4], p=immigration_probabilities[fly[2], :]) if fly[0] else fly[1] for fly in zip(flies.migrate, flies.release, flies.stock)]
    # This step zips together migrate, release and stock for each fly
    # if the fly migrates a new cage is chosen based on immigration _probabilites 
    # else, the capture cage is the same as the release cage
    

def mate_flies(flies):
    """ For every cage, check for the presence both sexes and compute phenotype-ratios by sex.
    Decide which flies can mate (both sexes present in the box). Pick a partner (based on phenotype ratio of 
    the opposite sex). Check whether or not they have the same phenotype. 

     Arguments
     ---------
        flies: a DataFrame containing the output from settle_flies()
    
    Returns
    -------
        modifies the first argument in-place
    """

    sr = flies[['sex', 'capture']].groupby(['capture']).mean() # percent of males 
    sex_ratios = np.zeros(5)
    for i in range(5): # This works around having a cage without flies
        try:
            sex_ratios[i] = sr.loc[i]
        except:
            sex_ratios[i] = 0

    mated = [0 if sex_ratios[x] in [0,1] else 1 for x in flies.capture]
    flies.mated = mated

    stock_ratios = flies[['sex', 'capture', 'stock']].groupby(['sex', 'capture']).mean()

    mate_prob = [stock_ratios.loc[(1-fly[2],fly[1])].values[0] if fly[0] else 0 for fly in zip(flies.mated, flies.capture, flies.sex)]
    mates = np.random.binomial(1, p=mate_prob, size=len(mate_prob))
    flies.mate=mates

    inbred = [1 if (fly[0]and (fly[1] == fly[2])) else 0 for fly in zip(flies.mated, flies.stock, flies.mate)]
    flies.inbred = inbred
 

def assortative_mating(flies):
    """ Compute (number of same-phenotype pairs)/(total number of pairs) of a fly population.

     Arguments
     ---------
        flies: a DataFrame containing the output mate_flies()
    
    Returns
    -------
        float scalar
    """

    inbreeding = sum(flies.inbred)/sum(flies.mated)   
    return inbreeding  


def run_simulation(pop=20, n=100, emigration_probabilities=observed_emigration, immigration_probabilities=observed_immigration):
    """ Run a simulation step by step and report summary statistics of the output.

     Arguments
     ---------
        pop: number of flies (it is rounded up to a multiple of 20 by generate_flies)

        n: number of replicates 

        emigration_probabilities: a 2x2 numpy array containing emigration probabilities by sex and stock

        immigration_probabilities: a 2x5 numpy array containing immigration probabilities by stock and cage
    
    Returns
    -------
        lisf of length n containing the assortative mating score for each population 

        print out the mean, standard deviation and standard error for each population 
    """

    print('Simulation started with %d flies and %d replicates' %(pop, n))
    print('    -----------------------------------------------')
    results = np.empty(n)

    for i in range(n):
        if i%100==0:
            print('ran %d times' %i) 

        a = generate_flies(pop)
        emigrate_flies(a, emigration_probabilities=emigration_probabilities)
        settle_flies(a, immigration_probabilities=immigration_probabilities)
        mate_flies(a)

        results[i] = assortative_mating(a)  # add this run's score to the list

    m = np.mean(results)
    d = np.std(results)
    e = d/(n**0.5)

    print('finished %d replicates' %(n))
    print('mean score: %1.3f, sd: %1.3f, se: %1.3f' %(m, d, e))
    print('')
    
    return (results)

np.random.seed(1802)

#Simulations

print('Observed emigration and immigration', )
observed_i = run_simulation(pop=200, n=1000)
print('')

print('Observed emigration, random immigration')
random_i = run_simulation(pop=200, n=1000, immigration_probabilities=random_immigration)
print('')

print('Observed emigration, deterministic immigration')
deterministic_i = run_simulation(pop=200, n=1000, immigration_probabilities=deterministic_immigration)
print('')

    
```
